# The radio-protective effects of (-)- Epigallocatechin-3-gallate (EGCG): regulating macrophage function in radiation-induced intestinal injury

**DOI:** 10.1101/2022.05.04.490595

**Authors:** Jia Gu, Zhibing Tang, Jiaxin Li, Yilin Cui, Shanghai Liu, Qing Guo, Yuzhong Chen, Sichen Lv, Lin Zhao, Jiaying Xu

**Affiliations:** State Key Laboratory of Radiation Medicine and Protection, School of Radiation Medicine and Protection, Collaborative Innovation Center of Radiation Medicine of Jiangsu Higher Education Institutions, Soochow University, Suzhou, China; Department of Orthopaedic Surgery, Suzhou Kowloon Hospital, Shanghai Jiaotong University School of Medicine, Suzhou, China; Yancheng Municipal Center for Disease Control and Prevention, Yancheng, China

## Abstract

Accidental and medical radiation exposure may cause radiation-induced intestinal injury (RIII), a catastrophic disease requiring efficient therapies. (-)-Epigallocatechin-3-gallate (EGCG), a major constituent of green tea, has been shown to have potent biological activity as well as strong anti-inflammation effects. Here, we demonstrate that EGCG treatment not only protects mice against total body irradiation (TBI)-induced toxicity and weight loss but also alleviates whole abdominal irradiation (WAI)-induced intestinal injury. EGCG promotes proliferation and survival of intestinal stem cells, inhibits radiation-induced apoptosis and inflammation. At the same time, EGCG preserves the composition of the gut microbiota in WAI mice. In vitro, we demonstrate that EGCG regulates the release of radiation-induced inflammatory factors by inhibiting inflammatory pathways such as toll-like receptor signaling pathway in peritoneal macrophages. Mechanistically, the radioprotective effect of EGCG was likely attributable to its preservation on macrophages and the colonized gut microbiota composition, thus relieving the intestinal inflammation. EGCG provides a novel strategy to mitigate RIII and improve the prognosis of patients after radiotherapy.

## Introduction

Due to the high radiosensitivity of intestine, radiation-induced intestinal injury (RIII) is the overwhelmingly most important barrier to patients receiving radiotherapy or exposed to accidental radiation (Harfouche et al., 2010; Pierre et al., 2016). RIII has symptoms such as nausea, diarrhea, bleeding, intestinal perforation and could even be fatal, which seriously impacts the survival condition of the patients (Chaves-Pérez et al., 2019; Ruiz-Tovar et al., 2009; Shadad et al., 2013). Previously, studies on the improvement and mechanism on herbal extractions with fewer side effects, simple extraction, and remarkable efficiency against RIII have received great attention. (-)-Epigallocatechin-3-gallate (EGCG), a major bioactive catechin derived from green tea (Stenvang et al., 2016), has numerous biological activities, such as anti-bacterial (Reygaert, 2018; Steinmann et al., 2013), anti-inflammatory (Wang et al., 2021; Zhong et al., 2012), and radioprotective activities (Kępka et al., 2019; Yi et al., 2020). EGCG has been demonstrated to ameliorate esophagitis (Li X et al., 2020; Zhu et al., 2021), oral mucositis (Zhu et al., 2020), and acute skin damage (Zhao et al., 2016; Zhu et al., 2016) induced by radiation in clinical trials, which suggests that it has a prospect in clinical application. Xie LW et al. (Xie et al., 2020) have investigated the effects of EGCG on RIII and revealed the key role of ROS and Nrf-2 in the radioprotection in intestinal epithelial cells. Moreover, Emami H et al. (Emami et al., 2014) found that green tea could relieve diarrhea and vomiting in patients undergoing pelvic or abdomen radiotherapy. Currently, the mechanism of EGCG alleviating RIII symptoms remains unknown.

Extensive studies of radioprotector focused on improving the radiation-induced destruction of intestinal epithelial cells (Li M et al., 2020; Qin et al., 2021; Zhang C et al., 2020). The mechanical barrier formed by intestinal epithelial cells, together with chemical, biological and immunologic barriers, forms the intestinal microenvironment and maintains the intestinal homeostasis (Beumer et al., 2021; Hageman et al., 2020; Luissint et al., 2016). However, little is known about the effect of radioprotector on the radiation-induced damage of immunologic barriers. Macrophage, as an important part of immunologic barriers, plays a central role in intestinal homeostasis and inflammation (Krause et al., 2015; Nighot et al., 2021). Studies have shown that macrophages are involved in tissue remodeling, removing dying cells (Grainger et al., 2017), avoiding the injury of excessive inflammation, maintaining the gut epithelial layer (Grainger et al., 2013) and so on. In addition, macrophages interact with gut microbiota, which is reflected in the critical role of gut microbiota shaped by macrophages (Earley et al., 2018), and the enhancing of antimicrobial activity induced by microbial metabolite (Schulthess et al., 2019). However, the specific role of macrophages that against RIII has not been studied; moreover, whether RIII can be relieved by the recovery of macrophages remains undocumented.

In the present study, we evaluated the therapeutic potential and molecular mechanisms of EGCG treatment in a lethal irradiation and a RIII model mice model respectively based on our previous study (Gu et al., 2020). We demonstrated that EGCG increases survival and prevents weight loss in mice after lethal irradiation. In RIII model, EGCG protected intestinal injury induced by radiation and retained the composition of gut microbiota. More importantly, EGCG inhibited the HMGB1/TLR4 inflammatory pathways in macrophages, which regulated the downstream inflammatory factors. Thus, our findings provide new insights into the function and underlying protective mechanism of EGCG against radiation-induced intestinal toxicity.

## Results

### EGCG increases survival and prevents weight loss in mice after TBI

To determine the radioprotective effects of EGCG in mice, we used 10 Gy total body irradiation (TBI) to cause lethal exposure and observed for following 14 days (Figure 1a). The results show that all the mice in the vehicle group and 2.5 mg/kg EGCG group had died, while half of mice in 10 mg/kg EGCG group remained alive at the 14^th^ day (Figure 1b). The average survival time of the vehicle, 2.5 mg/kg EGCG and 10 mg/kg EGCG was 6.6 days, 8.1 days and 11.2 days (*P*<0.05 vs vehicle; Figure 1c), respectively. Moreover, EGCG treatment significantly reduced body weight loss in the irradiated mice (Figure 1d), these results indicate that EGCG protects mice against lethal exposure induced mortality rate and weight reduction.

**Figure 1.**
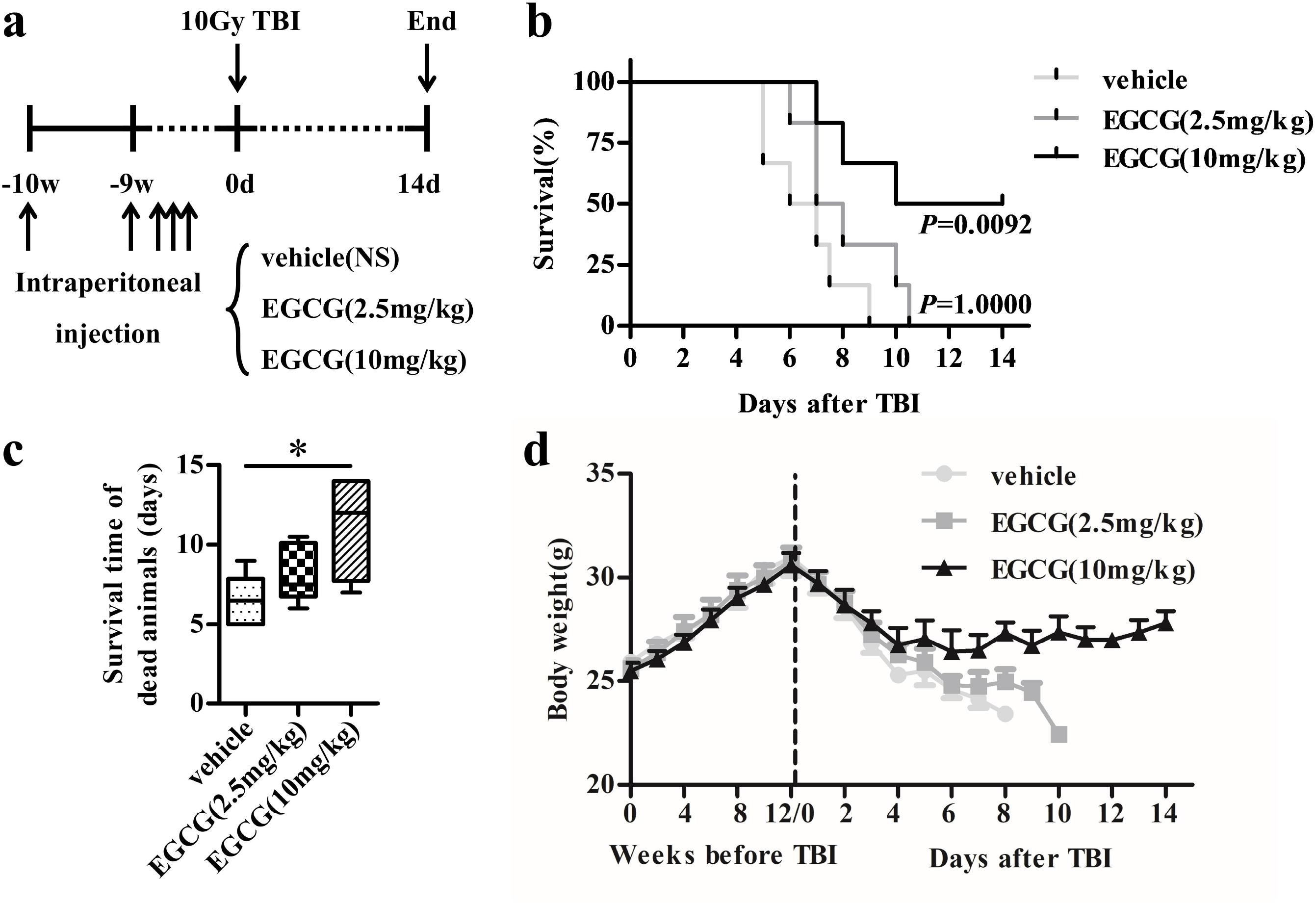
EGCG increases survival and prevents weight loss in lethally irradiated mice. (**a**) Mice received intraperitoneal injection of vehicle (0.9% normal saline), 2.5 or 10 mg/kg EGCG respectively for 10 consecutive weeks before TBI. All mice were exposed to a single dose of 10 Gy TBI. (**b**) Kaplan-Meier survival analysis of mice after exposure to 10Gy TBI. *P* = 1.0000 by log-rank testing between TBI-exposed mice treated with EGCG (2.5mg/kg)/vehicle, *P* = 0.0092 by log-rank testing between TBI-exposed mice treated with EGCG (10mg/kg)/vehicle, *n* = 6. (**c**) Mean survival time of C57BL/6J mice in three groups, *n* = 6, \**P* <0.05. The top and bottom boundaries of each box indicate the 75^th^ and 25^th^ quartile values, respectively, and lines within each box represent the 50^th^ quartile (median) values. Ends of whiskers mark the lowest and highest diversity values in each instance. (**d**) Body weights of mice in each group, *n* = 6.

### EGCG attenuates small intestinal injury in mice after WAI

Based on our previously established RIII model (Gu et al., 2020), we further explored the relationship between the protective effect of EGCG on intestinal injury and immune response in irradiated mice (Figure 2a). The results showed that the spleen and thymus indexes decreased significantly in WAI group, while EGCG+WAI group was improved especially at 3.5 days after WAI (Figure 2b, c). Blood routine analysis showed that the number of white blood cells (WBC) and monocytes (Mon) significantly decreased after WAI. EGCG pretreatment increased the number of WBC at 7 days after WAI (Figure 2d, Figure 2-table supplement 1), but there was no significant difference between WAI and EGCG+WAI group in the number of Mon (Figure 2e). As shown in Figure 2f, WAI caused significant visible acute intestinal injury such as poor elasticity and thinning of intestinal walls at 3.5 days after WAI, and intestinal swelling was apparent at 7 days after WAI. In the EGCG+WAI group, the intestinal tract was near to the normal state with only slight or no air accumulation. These phenomena were confirmed via H&E staining and PAS staining (Figure 2g, h), revealed reduction in villus height and the number of goblet cells at 3.5 days and 7 days following WAI (Figure 2i, j). In contrast, the crypt-villi architecture was well preserved in EGCG+WAI group.

**Figure 2.**
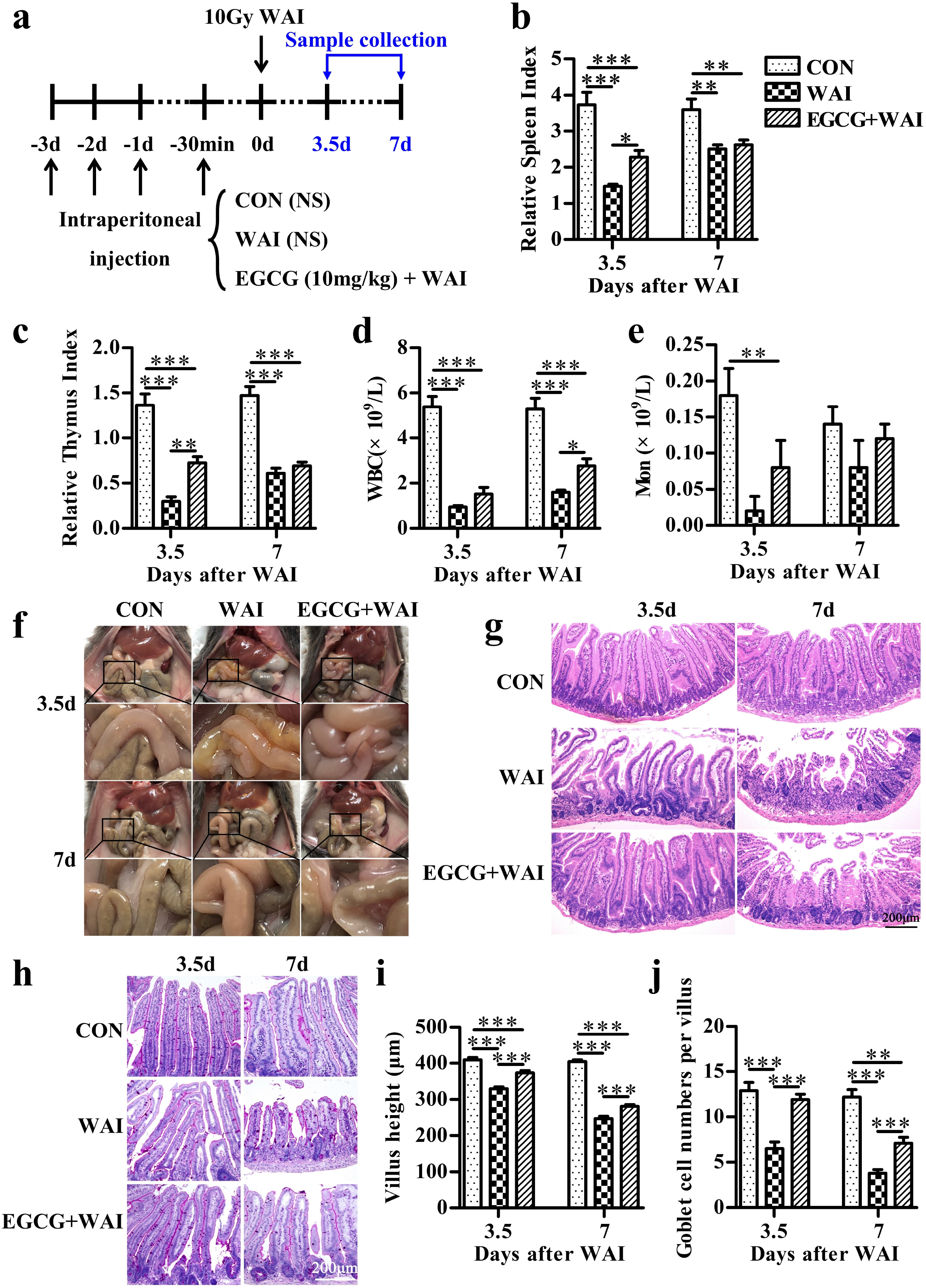
EGCG promotes small intestinal structure repair in nonlethally irradiated mice. (**a**) Mice were intraperitoneally injected with 5% chloral hydrate and received 10 Gy WAI. The mice were intraperitoneally injected with 10 mg/kg EGCG or an equivalent volume of saline for 3 consecutive days and 30 min before WAI. The spleen index (**b**) and thymus index (**c**) in the three groups at 3.5 days and 7 days after WAI. *n* = 6, \**P* < 0.05, \*\**P* < 0.01, \*\*\**P* < 0.001. Bar graph comparing the number of WBC (**d**) and Mon (**e**) in the three groups. *n* = 6, \**P* < 0.05, \*\**P* < 0.01, \*\*\**P* < 0.001. (**f**) Representative images of the respective intestinal tracts. The morphology of the jejunum of mice in the three groups at 3.5 days and 7 days after 10 Gy WAI was shown by H&E staining (**g**) and PAS staining (**h**) (Scale bar = 200 μm). Bar graph comparing villus height (**i**) and the number of goblet cells (**j**) in the three groups. *n* = 6, \*\**P* < 0.01, \*\*\**P* < 0.001.

### EGCG promotes proliferation and survival of intestinal stem cells, inhibits radiation-induced apoptosis and inflammation

We further evaluated the effects of EGCG on the levels of proliferation, differentiation, regeneration and apoptosis cells by IHC staining and TUNEL assay after WAI. Lgr5 is the marker of stem cell in the small intestine (Barker et al., 2007). Paneth cells, located at the bottom of the small intestinal crypts, limit bacterial invasion by secreting antimicrobial proteins, including lysozyme (Bel et al., 2017; Yu et al., 2020). Ki67 is a marker of regeneration of the epithelial layer (Figure 3a). Furthermore, TUNEL assay was applied to analyze apoptosis in the small intestinal tissues (Figure 3b). Compared to the CON group, the numbers of Lgr5^+^ cells per crypt, lysozyme^+^ crypts per circumference and Ki67^+^ cells per crypt decreased significantly in the WAI group and were higher in the EGCG+WAI group (Figure 3c-e). There were a large number of TUNEL^+^ cells in the crypts and villi in WAI group compared to the CON group, and EGCG+WAI group significantly decreased the apoptosis levels in the small intestine (Figure 3f). Alkaline phosphatase (ALP) is an endogenous enzyme and plays an important role in maintaining the integrity of intestinal epithelial cells (He et al., 2021; Sheng et al., 2021). At 3.5 days after WAI, ALP was lost from the jejunal (Figure 3g), the permeability of epithelial cells increased, causing increasingly expression of inflammatory factors (Figure 3h-j) (Lallès, 2019). EGCG treatment maintained the ALP activity of mouse jejunal epithelial cells and reduced the release of inflammatory factors including IL-6, TNF-α and IFN-γ. Taken together, these data demonstrate that EGCG treatment increases the expression of Lgr5, Ki67 and lysozyme, decreases the rate of apoptosis and reduces the level of inflammation in the small intestines of WAI-exposed mice.

**Figure 3.**
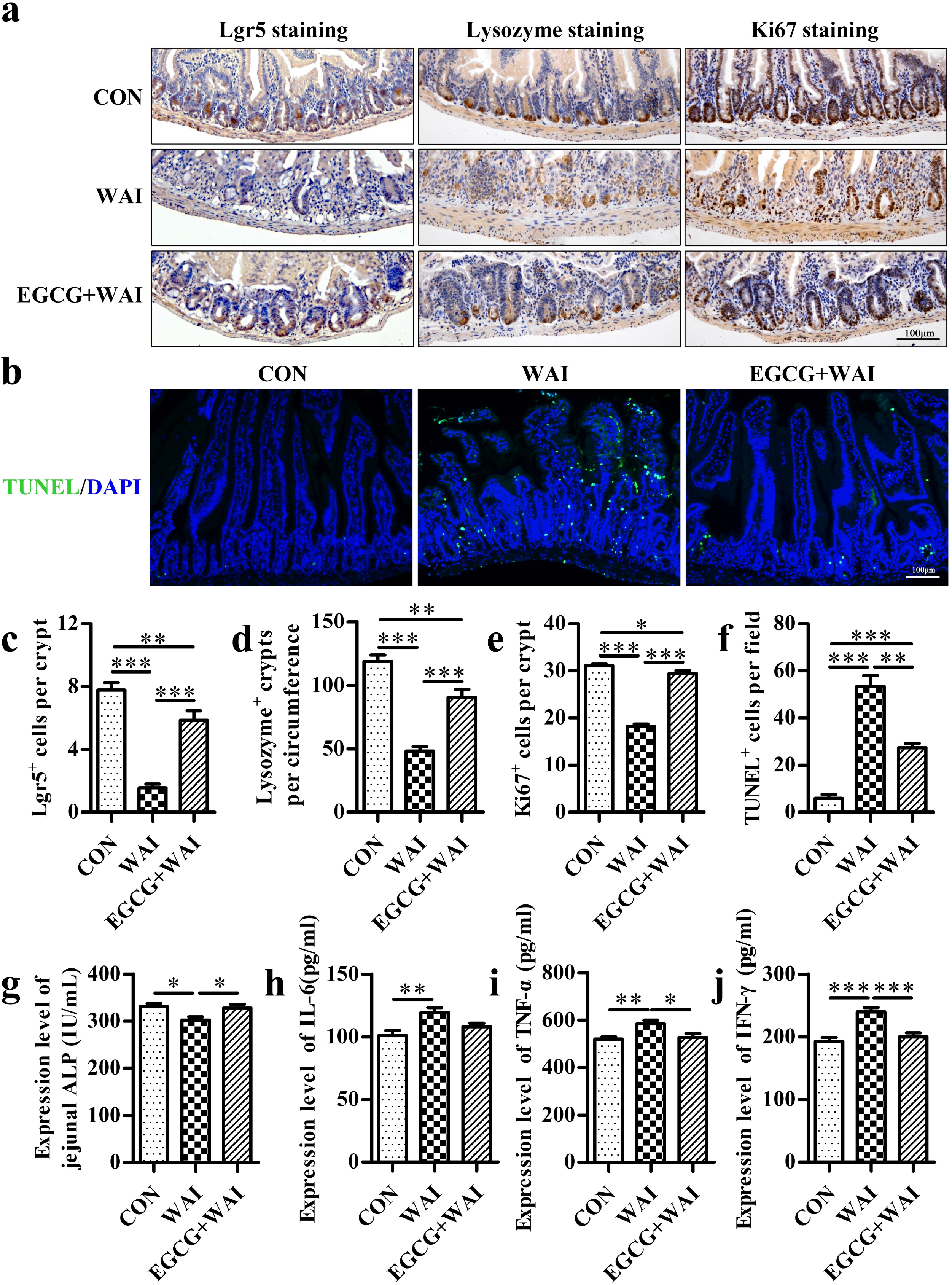
EGCG promotes crypt epithelial cell proliferation, decreases the rate of apoptosis and reduces the level of inflammation after WAI. (**a**) Representative images of the expression of Lgr5, lysozyme and Ki67 in cross-sections of the small intestine visualized by IHC staining at 7 days after WAI (Scale bar = 100 μm). (**b**) The frequency of apoptosis in the small intestines was assayed by TUNEL staining at 7 days after WAI (Scale bar = 100 μm). Histograms showing number of Lgr5^+^ cells per crypt (**c**), lysozyme^+^ crypts per circumference (**d**) and Ki67^+^ cells per crypt (**e**) determined from panel (a). *n* = 6, \**P* < 0.05, \*\**P* < 0.01, \*\*\**P* < 0.001. (**f**) Histograms showing number of TUNEL^+^ cells (green) determined from panel (b). *n* = 6, \*\**p* < 0.01, \*\*\**P* < 0.001. (**g**) Histograms showing expression level of jejunal ALP at 3.5 days after WAI via ELISA technique. *n*= 5, \**P* < 0.05. Histograms showing expression level of serum inflammatory factors including IL-6 (**h**), TNF-α (**i**) and IFN-γ (**j**) at 3.5 days after WAI via ELISA technique. *n* = 5, \**P* < 0.05, \*\**P* < 0.01, \*\*\**P* < 0.001.

### EGCG alters the composition of the gut microbiota in WAI mice

Gut microbial dysbiosis has been reported to affect the radiosensitivity of hosts (Cui et al., 2017; Xiao et al., 2018). We therefore performed 16S rDNA high-throughput sequencing to study the effects of EGCG on bacterial taxonomic composition after WAI (Figure 4-table supplement 1). The Chao 1 index, the observed species number and the Shannon index analysis exhibited no significant difference of α-diversity among the three groups (Figure 4a), indicating that 10 Gy WAI unaltered the species of gut microbiota. Principal coordinate analysis (PCA) showed gut microbiota structures were substantially changed after WAI exposure and EGCG treatment (Figure 4b). We further applied linear discriminant analysis (LDA) effect size (LEfSe) analysis to identify the differences in taxa abundance between the three groups (Figure 4c, d). Combined with the bar chart results, we found that the relative abundance of *Escherichia_Shigella* and *Bacteroides* were higher while the relative abundance of *Muribaculaceae* and *Duncaniella* were lower in the WAI group than in the CON group at the genus level; however, treatment with EGCG restored these changes (Figure 4e, f). These data demonstrate that EGCG does not affect the abundance but restores the structures of gut microbiota in WAI mice.

**Figure 4.**
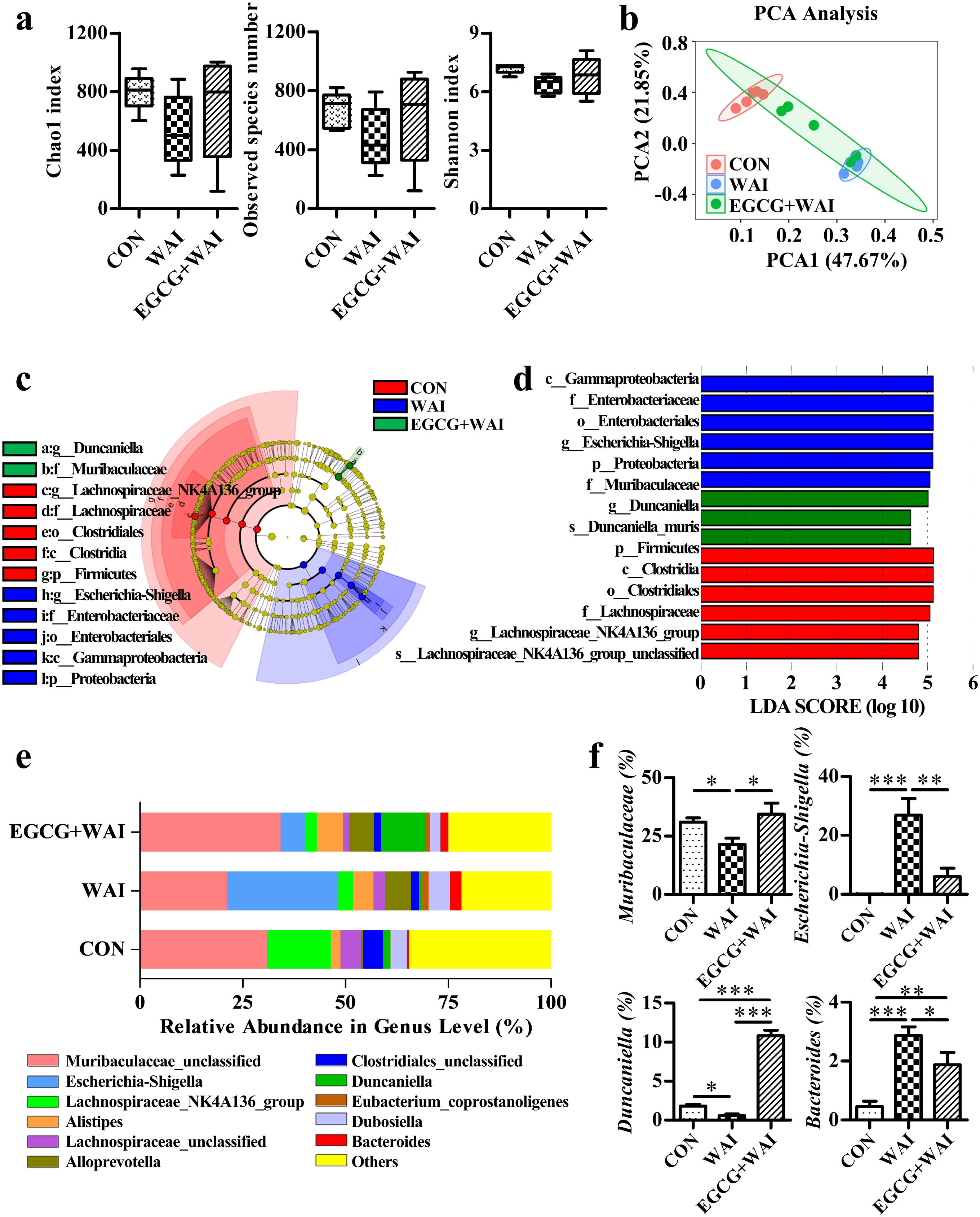
EGCG reverses the alterations in gut microbiota induced by WAI. (**a**) Box-plots showing the alpha diversity of gut microbiota, including the Chao1 index, the observed species number and the Shannon index. *n* = 5, *P* > 0.05. The top and bottom boundaries of each box indicate the 75^th^ and 25^th^ quartile values, respectively, and lines within each box represent the 50^th^ quartile (median) values. Ends of whiskers mark the lowest and highest diversity values in each instance. (**b**) Cluster analysis by using PCA. The first two principal components (PC1 and PC2) from PCA are plotted for each sample. The percentage variation covered in the plotted principal components is marked on the axes. Each spot represents one sample, and each group of mice is labeled by a different color, *n* = 5. (**c**) Cladogram derived from LEfSe analysis of 16S sequences. Regions in red, blue, green indicate clades that were enriched in CON, WAI, EGCG+WAI group respectively. Regions in yellow indicate clades with no significant difference. From inside to outside, cladogram denotes the taxonomic level of phylum, class, order, family and genus. (**d**) LEfSe results showed that bacteria abundances were significantly different between the three groups, and indicated the effect size of the most differentially abundant bacterial taxon in the small intestine, *n* = 5. The alpha value for the factorial Kruskal-Wallis test among classes <0.05 was considered significant and the threshold on the logarithmic LDA score for discriminative features was 4.5. (**e**) The relative abundance of the top 11 gut microbiota at the genus level in mice, *n* = 5. (**f**) Histograms showing the proportion of gut microbiota with significant differences in genus level. *n* = 5, \**P* < 0.05, \*\**P* < 0.01, \*\*\**P* < 0.001.

### EGCG recovers the viability of macrophages after radiation

To determine the protective effect of EGCG against RIII, we used peritoneal macrophages and RAW264.7 macrophages as in vitro model systems. First, we performed CCK-8 assays to evaluate cell viability. EGCG treatment at concentrations higher than 50 μM significantly retained the growth of both macrophages (Figure 5a). Thus, EGCG at lower concentrations of 2.5, 5, 10, 25 μM was used to determine the radioprotective effects. Treatment with EGCG offered significant protection against loss of cell viability induced by radiation. Considering the radioprotective effect after 24 h culture in macrophages, we used EGCG at a concentration of 10 μM (Figure 5b, c). The results of live/dead assays were consistent with those of CCK-8 assays (Figure 5d). These data suggest that EGCG attenuates damage induced by radiation in both macrophages.

**Figure 5.**
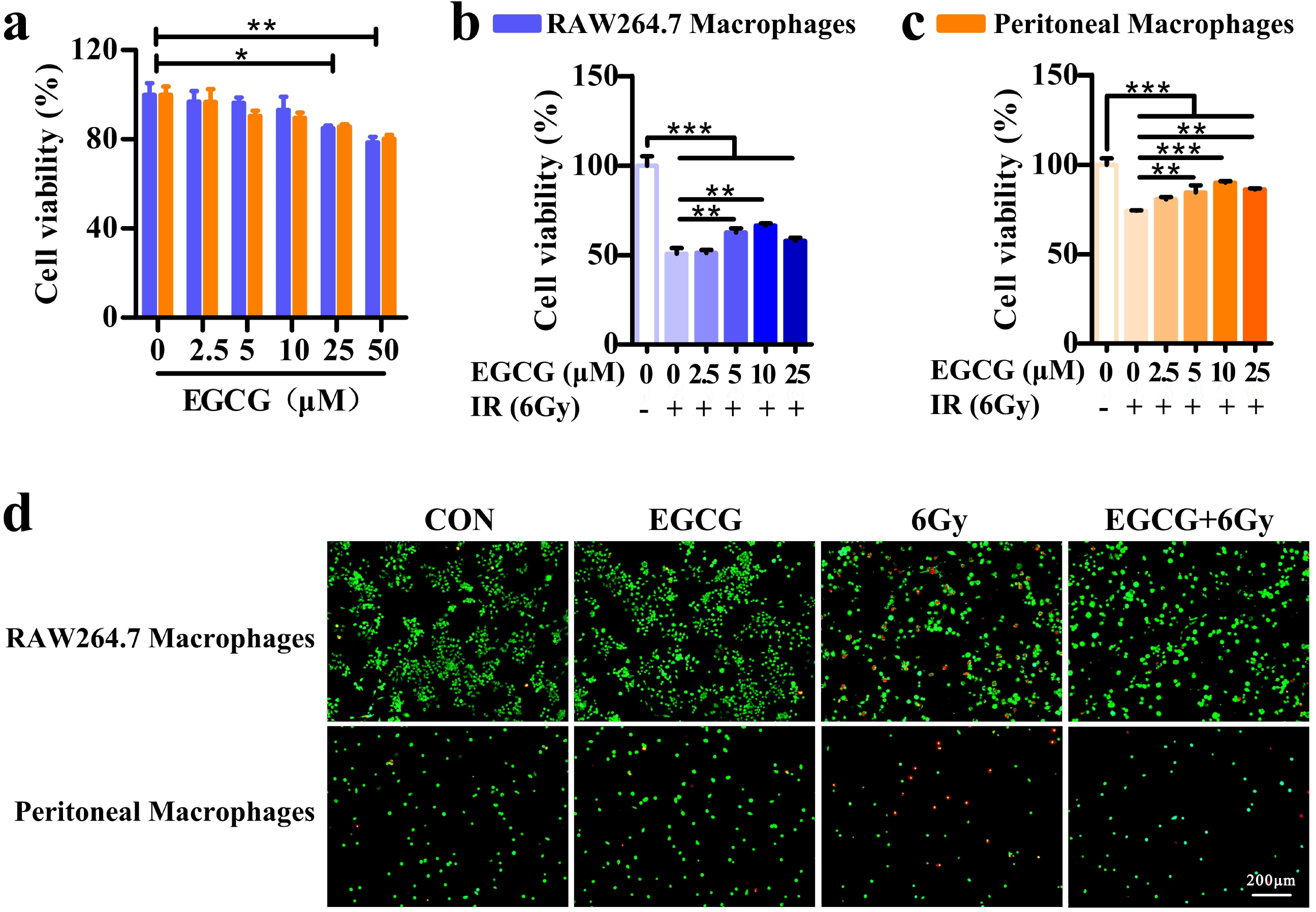
EGCG inhibits radiation-induced suppression of viability in cells. (**a**) Cytotoxicity of EGCG in peritoneal macrophages and RAW264.7 macrophages. Cells were treated with the indicated concentrations of EGCG for 24 h. Cell viability was monitored using the CCK-8 assay. *n* = 5, \**P* < 0.05, \*\**P* < 0.01. (**b**) Effects of EGCG on cell viability post radiation in RAW264.7 macrophages. RAW264.7 macrophages were sham-irradiated or irradiated with 6 Gy irradiation 30 min after receiving indicated concentrations of EGCG. Cells were cultured for 24 h and cell viability was monitored using the CCK-8 assay. *n* = 5, \*\**P* < 0.01, \*\*\**P* < 0.001. (**c**) As above, peritoneal macrophages were cultured for 24 h and cell viability was monitored using the CCK-8 assay. *n* = 5, \*\**P* < 0.01, \*\*\**P* < 0.001. (**d**) Live/Dead assay of EGCG and 6 Gy treatment on macrophages. Cells were sham-irradiated or irradiated with 6 Gy irradiation 30 min after receiving 0, 10 μM EGCG. A live (cells illuminated by green light)/dead (cells illuminated by red light) assay after 24 h treatments (Scale bar = 200 μm).

### EGCG attenuates radiation-induced inflammation in macrophages

To determine the mechanism underlying the protective effects of EGCG against radiation-induced macrophage activity, we examined the expression of 40 inflammatory factors in culture medium of irradiated macrophages (Figure 6a, and all data of proteins were provided in Figure 6-figure supplement 1). Combined with analysis by volcano plot shows that compared to EGCG+6Gy group, several proteins were significantly upregulated in the 6 Gy group, including Eotaxin-2, IL-6, TNF-α and BLC, while Eotaxin, IL-7, IL-17 and FP4 was significantly downregulated (fold-change > 1.5 or < −1.5, *P* < 0.05) (Figure 6b). KEGG enrichment analyses were further performed to investigate the enriched pathways of differentially expressed proteins in peritoneal macrophages. KEGG enrichment analysis revealed that differentially expressed proteins were significantly enriched in 20 pathways (Figure 6c, Figure 6-table supplement 1). According to the results, we verify the toll-like receptor signaling pathway via Western Blot. As expected, EGCG treatment inhibited the increase of HMGB1, NF-κB and TLR4 protein expression induced by radiation (Figure 6d, Figure 6-figure supplement 2). In summary, EGCG regulates the release of inflammatory factors such as IL-6 by inhibiting the inflammatory pathways of macrophages after irradiation.

**Figure 6.**
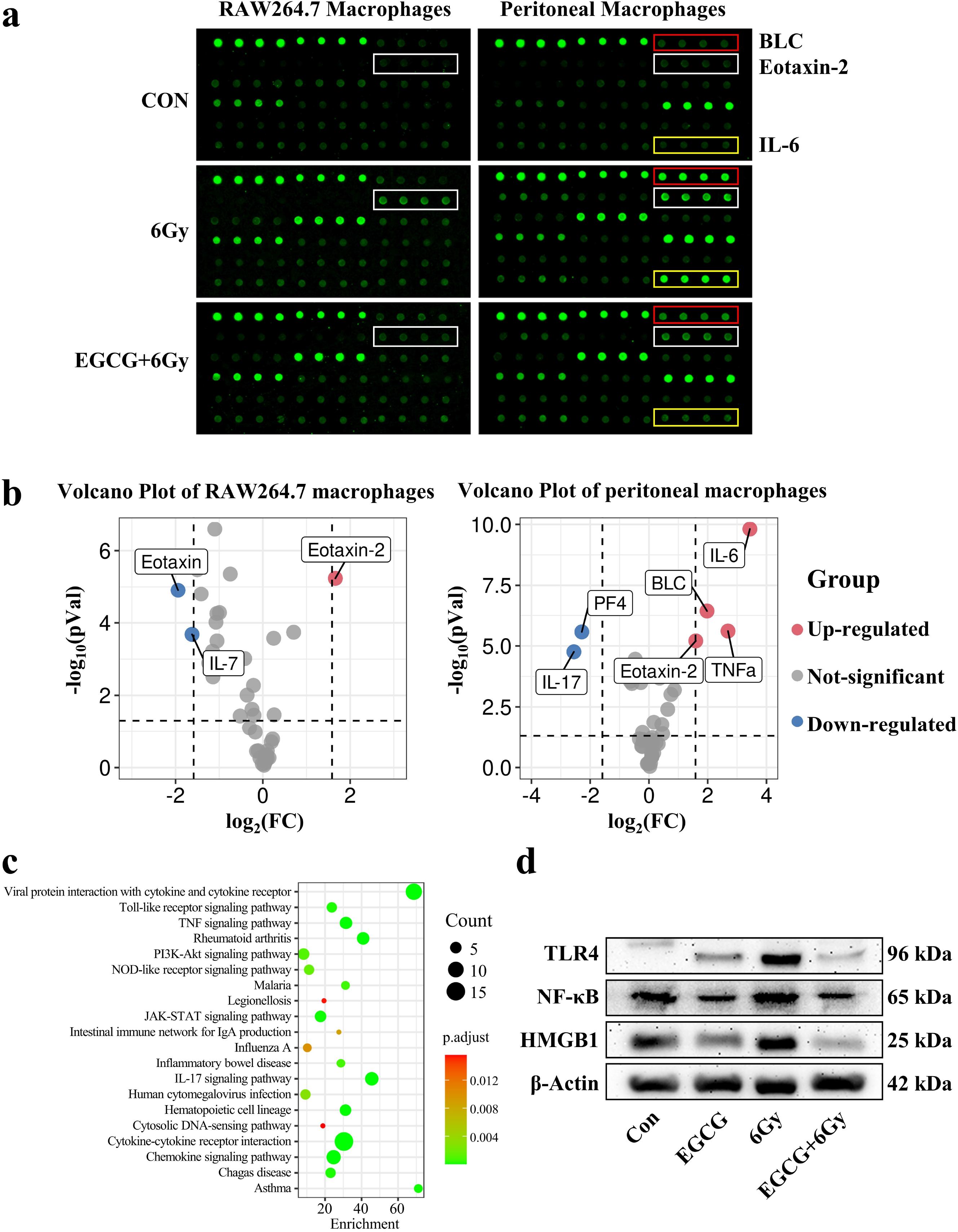
EGCG relieves inflammation induced by radiation in peritoneal macrophages. (**a**) Cells were sham-irradiated or irradiated with 6 Gy irradiation 30 min after receiving 0, 10 μM EGCG. The culture medium was collected after 24 h and concentrated. Representative results from the mouse inflammation array GS1.16 mouse inflammatory proteins were profiled in culture medium from macrophages using the mouse inflammation array GS1. (**b**) Volcano plot of the protein expression profile in mouse inflammation array GS1. The log2 (Fold Change) value of the differential protein expression is taken as the abscissa, and the negative logarithm-log10 (P Value) of the significance test P values between the two groups is taken as the ordinate. Red represents upregulation, blue represents downregulation, and gray represents no difference. Blue and red spots represent up-and downregulated > 1.5-fold or < −1.5-fold and P < 0.05 in 6 Gy versus EGCG+6Gy group. (**c**) The enriched KEGG pathways of the differentially expressed proteins in peritoneal macrophages between 6Gy group and EGCG+6Gy group, according to *P* < 0.05. (**d**) Proteins of peritoneal macrophages treated with 0, 10 μM EGCG 30 min before sham-irradiated or 6 Gy irradiation were extracted and Western blot was performed to detect the protein level of HMGB1, NF-κB and TLR4. β-actin was used as a loading control.

## Discussion

EGCG has been reported to be a remarkable molecule belonging to green tea catechin with multiple health benefits and low toxicity (Chakrawarti et al., 2016). In vitro and animal models, the use of EGCG has been shown to inhibit a high variety of cancer processes (Singh et al., 2011). In vivo studies, EGCG acts as a radiosensitizer to synergize the treatment of cancer (Lecumberri et al., 2013), and also be able to ameliorates the adverse side effects derived from cancer therapy (Li X et al., 2020; Zhao et al., 2016; Zhu et al., 2016; Zhu et al., 2020; Zhu et al., 2021). Previous studies have revealed that treatment with EGCG protects rodents from intestinal damage after TBI up to 9 Gy (Emami et al., 2014). On the basis of previous studies, we speculated that EGCG might function as a potential therapeutic radioprotectant to treat RIII, and further explored the underlying mechanism from a novel immunity perspective.

We first validated that EGCG treatment improved overall survival following 10Gy TBI. Then we determined whether EGCG might ameliorate RIII in a WAI murine model (Gu et al., 2020). The results indicated that EGCG retained the normal crypt-villus architecture of jejunal tissue following 10 Gy WAI. Due to the vigorous proliferation of intestinal stem cells, which are composed of crypt-base columnar cells intermingling with Paneth cells (Barker et al., 2007), the small intestine is highly sensitive to radiation (Harfouche et al., 2010; Potten et al., 2004). The mechanism of high radiosensitivity may be pro-apoptotic protein accumulation (Li et al., 2019; Zhou et al., 2015). In addition, ALP is an enzyme secreted into the intestinal lumen by enterocytes, maintains gut homeostasis (Lassenius et al., 2017) via reducing the release of proinflammatory cytokines (Riggle et al., 2013; Sheng et al., 2021) and the occurrence of bacterial translocation (Fawley et al., 2017; Plaeke et al., 2020). We found that EGCG treatment maintained the regenerative capacity of jejunal cells by augmenting Ki67^+^ cells, lysozyme^+^ Paneth cells and declining TUNEL^+^ cells in WAI mice. Treatment with EGCG increased the level of ALP expression and decreased the release of proinflammatory cytokines in WAI mice. Therefore, we suspected that EGCG treatment attenuates RIII by relieving excessive inflammation and maintaining the regenerative capacity of the intestinal epithelium.

Currently, the pathogenesis of RIII remains unknown, but it is believed that the combined effects of epithelial injury, vascular changes, microbial and immune factors lead to severe radiation enteropathy (Hauer-Jensen et al., 2014; Li et al., 2021). The gut microbiota plays an important role in maintaining intestinal barrier function, regulating immune homeostasis and protecting intestinal repair mechanisms (Ahlawat et al., 2021; Gasaly et al., 2021; Touchefeu et al., 2014). Extensive evidence already showed that gut microbiota may be a key driver of radiation enteropathy process (Gerassy-Vainberg et al., 2018; Reis Ferreira et al., 2019; Sokol et al., 2018). Wang et al. (Wang et al., 2019) used bacterial-epithelial co-cultures and found that compared with control microbiota, radiation enteropathy-derived microbiota induced epithelial inflammation and barrier dysfunction by enhancing TNF-α and IL-1β expression. We therefore employed 16S rDNA gene sequencing and found that dysbiosis was observed in mice after WAI, which was characterized by significantly higher abundance of *Escherichia_Shigella* and *Bacteroides* and lower abundance of *Muribaculaceae* and *Duncaniella*. Treatment with EGCG restored the composition of gut microbiota in WAI mice by reducing the proportion of harmful bacteria such as *Escherichia_Shigella.*

Interestingly, recent studies indicated that *Shigella* deploy multiple mechanisms to stimulate inflammasomes (Mitchell et al., 2020; Sanchez-Garrido et al., 2020), thereby inducing rapid macrophage cell death (Suzuki et al., 2014). The intestinal tract is the largest independent immune system in the body (Doe, 1972), in which macrophages play a crucial role (Han et al., 2021; Krause et al., 2015; Nighot et al., 2021). A proportion of the macrophages are derived from the migration and differentiation of Ly6C^hi^ blood monocytes (Bain et al., 2014), and the others are self-maintaining gut-resident macrophages with longevity (De Schepper et al., 2019). Previous evidence from murine studies suggested that macrophages are capable to clear the bacteria from the tissue and secrete cytokines to establish the local homeostatic immune cell network (Gross et al., 2015; Niess et al., 2005; Vallon-Eberhard et al., 2006). What’s more, after radiation, macrophages sense inflammatory molecules through the expression of toll-like receptor 4 (TLR4) for high mobility group box 1 (HMGB1) and the downstream signaling leads to release of pro-inflammatory cytokines that initiate the inflammatory cascade (Kim et al., 2017; Shi et al., 2018). In this study, our results suggest that EGCG are capable to protect cell viability of macrophages after radiation, regulate the inflammatory signaling pathway (e.g., toll-like receptor signaling pathway) and the release of inflammatory factors (e.g., Eotaxin-2, Eotaxin, IL-6, TNF-α, etc.) of peritoneal macrophages. Overall, we speculate that EGCG protects mice intestinal system from radiation induced injury by a direct mechanism involved by intestinal stem cells, and an indirect mechanism involved by macrophage, which regulated by gut microbiota (Figure 7).

**Figure 7.**
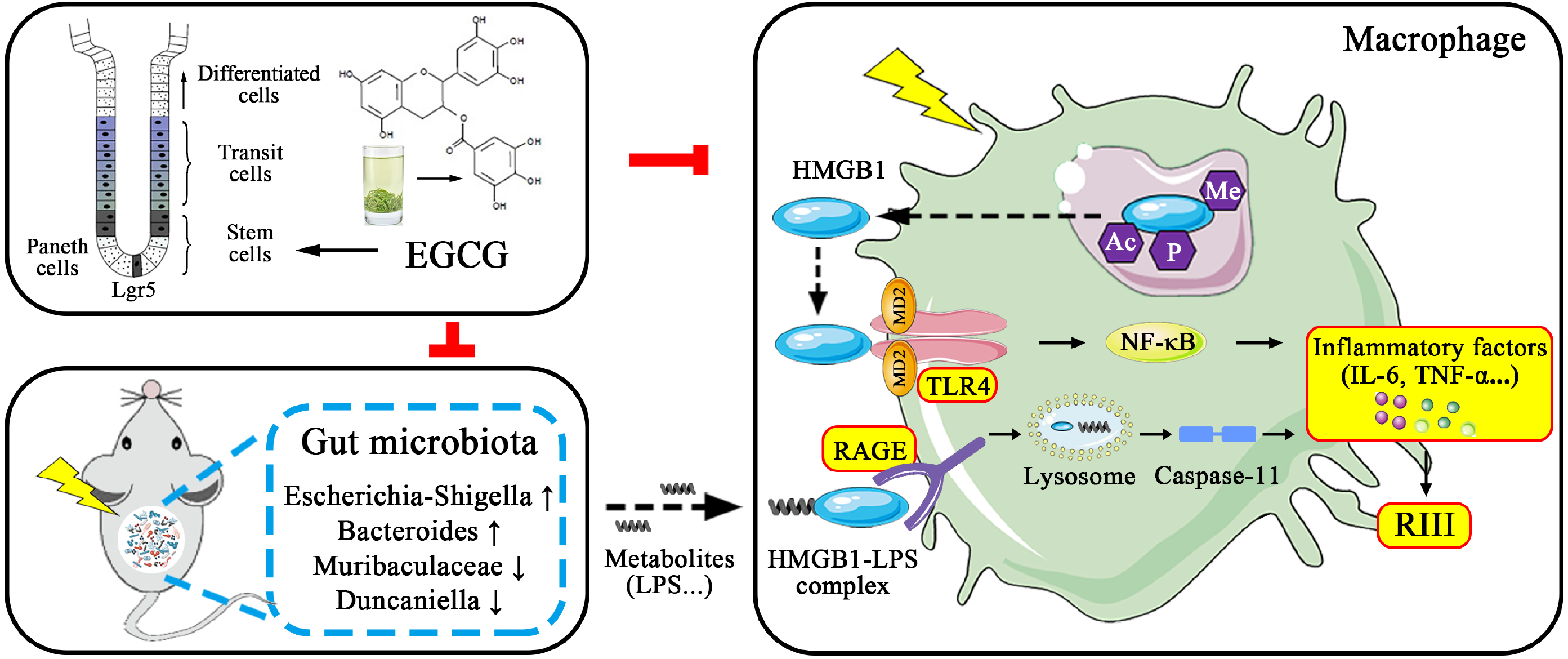
The underlying mechanism of the radio-protective effect of EGCG.

In summary, the findings presented in this study show that EGCG can offer substantial protection against RIII in WAI mice by protecting intestinal immune microenvironment. Thus, we provide a new direction for exploring the effects of radiation protection drugs.

## Materials and methods

### Animals

Six-to eight-week-old male C57BL/6J mice were obtained from Shanghai Laboratory Animal Center of Chinese Academy of Science (Shanghai, China). All the mice were housed in a polycarbonate cage with a controlled environment (22 °C±2 °C, 60%±5% relative humidity, and 12 h light/dark cycle) and provided with standard mouse feed ad libitum throughout the experiment. All male mice in this study were of a pure C57BL/6J genetic background and separated into groups randomly. All experimental procedures were performed in accordance with the Guidelines in the Care and Use of Animals and with the approval of Soochow University Animal Welfare Committee.

### Irradiation and EGCG treatment

An X-RAD 320 Irradiator (Precision X-ray Inc., USA) at a dose rate of 2.0 Gy per minute was used for mice experiments. For the survival experiments, mice were randomly divided into three groups as follows: (1) vehicle; (2) 2.5 mg/kg EGCG; (3) 10 mg/kg EGCG (Yang et al., 2021). For the remaining experiments, the mice were randomly divided into three groups as follows: (1) control (CON); (2) whole abdominal irradiation + vehicle (WAI); (3) EGCG + WAI. WAI was delivered using a lead shielding so that whole abdominal was irradiated and the other parts of the mouse were shielded. Control mice were sham-irradiated. EGCG was purchased from Sigma-Aldrich (St Louis, MO, USA). For the animal experiments, EGCG was dissolved in 0.9% normal saline (NS) with a stock concentration of 5 mg/mL. For the cell experiments, EGCG was dissolved in phosphate buffered saline (PBS) with a stock concentration of 5 mg/mL. The doses and schedules for administration of EGCG and radiation are provided in the relevant figure legends.

### Histopathological studies

The small intestine samples were harvested and fixed in 10% buffered formaldehyde– saline solution. Then, 4 μm sections of paraffin-embedded samples were stained with hematoxylin and eosin (H&E, Sigma-Aldrich), periodic acid-Schiff (PAS, Servicebio) and immune-histochemical reagents (IHC) as previously described (Gu et al., 2022). For IHC staining, deparaffinized sections were rehydrated and stained using specific antibodies against Leucine-rich repeat-containing G-protein-coupled receptor 5 (Lgr5, ab219107, Abcam, USA), Lysozyme (ab76784, Abcam, USA) or Ki67 (ab15580, Abcam, USA), then stained with biotinylated secondary antibodies. Detection was performed with an avidin-biotin horseradish peroxidase complex, using 3, 3’-diaminobenzidine (Servicebio) as the chromogen. The samples were photographed under a microscope (DMi8, Leica, Germany) and semi-quantitatively scored by a board-certified pathologist who was blinded to experimental conditions.

### Terminal deoxynucleotidyl transferase dUTP nick end labeling (TUNEL) assay

The 4 μm-thick intestinal sections were treated according to the manufacturer’s protocols (Servicebio).

### Blood analysis

Collected blood samples were used for routine blood test (BC-2800Vet, Mindray, China) to determine the levels of white blood cell (WBC), number of lymphocyte (Lymph), monocyte (Mon), granulocyte (Gran), red blood cell (RBC), hemoglobin (HGB), and mean corpuscular hemoglobin concentration (MCHC). The remaining blood samples were immediately centrifuged at 3000 g for 10 minutes at 4 °C. The serum was collected and stored at −80 °C for measurement of ELISA.

### Enzyme-linked immunosorbent assay (ELISA)

The obtained jejunal samples were homogenized with a tissue homogenizer at 4 °C for 10 min and then centrifuged at 3000 rpm for 10 min. The supernatants were stored at −80 °C. Serum inflammatory factors interleukin-6 (IL-6), tumor necrosis factor-α (TNF-α), interferon-γ (IFN-γ) and mouse alkaline phosphatase (ALP) in the supernatants were tested according to kit instructions using quantitative sandwich assay (JYMBio, China) to determine the concentrations. OD value was measured by multimode plate reader (Synergy NEO, BioTek, USA) at 450 nm, and the levels of each factor were calculated.

### Bacterial diversity analysis

DNA extraction and 16S ribosomal DNA (rDNA) gene amplification were performed as described previously (Gu et al., 2022). The 16S rDNA V3-V4 gene was analyzed to evaluate the bacterial diversity using an Illumina MiSeq technology (Lc-Bio Technologies Co., Ltd., China). All data of microbiota determined by 16S rDNA are provided in Figure 4-table supplement 1.

### Cell culture and culture conditions

RAW264.7 cell line and small intestinal epithelial cell line-6 (IEC-6) were purchased from Cell Resource Center, Shanghai Institute for Biological Sciences (SIBS, China). RAW264.7 macrophages were cultured in Dulbecco’s modified Eagle’s medium (DMEM, HyClone, USA) supplemented with 10% fetal bovine serum (FBS, Gibco, USA). IEC-6 cells were cultured in RPMI 1640 medium containing 10%FBS and 1%penicillin-streptomycin solution (Beyotime, China). Cells were maintained in a humidified incubator with 5% CO_2_ at 37 °C. The cells were exposed to 6 Gy radiation from the X-ray tube (Rad Source Technologies, USA) at a fixed dose rate of 1.15 Gy/min.

### Peritoneal macrophage isolation

Peritoneal macrophages were isolated from C57BL/6J mice as previously described with some modifications (Zhang N et al., 2020). Briefly, 30 min after intraperitoneal injection of 5 mL DMEM with 75% FBS in C57BL/6 mice, peritoneal macrophages were harvested from the peritoneal cavity with 5 mL of DMEM and centrifuged. The cells were resuspended with 10% FBS DMEM and cultured. After 2 h of cell culture, the unadherent cells were discarded and petri dish was washed twice, the obtained adherent cells were macrophages.

### Cell viability

Viability of cells was assessed by CCK-8 assay. Briefly, cells (5×10^3^ cells / well) were plated in 96-well plates, cultured for 24 h, and then treated with 0, 2.5, 5, 10, 25, or 50 μM EGCG for 30 min, followed by 0 or 6 Gy X-ray radiation. After 24 h treatment, CCK-8 (Beyotime, China) was added to each well (10 μL) for incubation for 1 h. Absorbance of each well was determined at 450 nm using a microplate reader (Synergy NEO, BioTek, USA). The experiment was independently repeated at least three times.

### Live/Dead assay

At the same time, cell viability was also tested using a Live/Dead assay. Cells were incubated in the absence or presence of 10 μM EGCG for 1 h before received 6Gy radiation or not. Biefly, 1 mL of PBS containing 4 μL of 2 mM ethidium homodimer-1 (EthD-1) assay solution and 2 μL of 50 μM calcein AM assay solution was prepared. Then, 200 μL of the Live/Dead solution was added to each well for 15 min in an incubator at 37°C. The staining solution was removed and the samples were then imaged under a fluorescence microscope (DMi8, Leica, Germany) with 494 nm (green, calcein) and 528 nm (red, EthD-1) excitation filters.

### Microarray data and data preprocessing

Microarray data and data preprocessing culture medium and cells obtained from the RAW264.7 macrophages and peritoneal macrophages was used to identify genes showing differential expression within culture medium and cells by Mouse Inflammation Array GS1 (GSM-INF-1-1, RayBiotech Company, USA). In this study, we used 40 cytokine-specific antibodies for detection, and each capture antibody was printed in quadruplicate on the glass. First, cell samples were lysed by cell lysis buffer containing protease inhibitor cocktail for 1 h on ice and centrifuged at 13, 000 rpm for 20 min at 4 °C. Then supernatant and culture medium samples were diluted (1:2) and added to each well to incubate with capture antibodies overnight. After washing, the arrays were incubated with a biotinylated antibody cocktail for 2 h at room temperature. After washing, Cy3 equivalent dye-streptavidin was added, and the fluorescent signal could be visualized using InnoScan 300 Microarray Scanner (Innopsys, France) equipped with a Cy3 wave length (green channel). The data were normalized using the RayBiotech analysis tool, an array-specific, Excel-based program that performs sophisticated data analysis on the raw numerical data extracted from the array scan. More details:

#### Microarray data and data preprocessing

Similar to a traditional sandwich-based ELISA, this array is a glass slide-based antibody array that can be used to conduct rapid, accurate expression profiling of hundreds of cytokines, chemokines, growth factors, proteases, soluble receptors and other proteins from any biological fluid.

#### Gene ontology and pathway enrichment analyses

To better understand the functions of the DEGs, Gene ontology (GO) enrichment and Kyoto Encyclopedia of Genes and Genomes (KEGG) pathway analyses were performed to discover the functions in which the DEGs participated. Functional enrichment analyses were based on Fisher’s exact test in the cluster Profiler package of R/Bioconductor, and the threshold values were set as count ≥ 2 and P < 0.05.

### Western blot

Protein level was analyzed using Western blotting analysis as described previously (Gu et al., 2022). Briefly, total protein lysates were obtained from peritoneal macrophages using RIPA (Beyotime, China) buffer containing protease inhibitors at 4 °C for 30 min and centrifuged at 12, 000 g for 15min at 4 °C. Antibody to high mobility group box 1 (anti-HMGB1, 66525-1-Ig) and toll-like receptor 4 (anti-TLR4, 19811-1-AP) were purchased from Proteintech. Antibody to nuclear factor kappa B p65 (anti-NF-κB, PB0073) and rabbit or mouse secondary antibodies were purchased from Boster Bio. The loading control was an anti-β-actin antibody (BM3873, Boster Bio). The membranes were developed by chemiluminescence using Superstar ECL Western blot substrate reagents (Beyotime). The bands were analyzed with Image J software (Version 1.8).

### Statistical Analysis

Each experiment was repeated at least three times. Data were expressed as the means ± standard error of the mean (s.e.m). The survival rates were analyzed using Kaplan-Meier survival testing. The significance of differences between two groups was evaluated using Student’s *t*-test. For comparing multiple groups, the differences were analyzed by one-way ANOVA (SPSS 24.0), followed by Fisher’s least significant difference (LSD) tests. Differences were considered statistically significant at *P* < 0.05.

## Data and code availability

The raw 16s rDNA sequencing data have been deposited in the GenBank Sequence Read Archive with the BioProject ID PRJNA833951.

## Acknowledgment

This work was supported by grants from the National Natural Science Foundation of China (No.81973024, 82073482), the Natural Science Foundation of the Jiangsu Higher Education Institutions of China (18KJA310006, 21KJB310006), the Open Project of Jiangsu Provincial Key Laboratory of Radiation Medicine and Protection (KJS2101), and cultivation and scientific research project of Suzhou Kowloon Hospital (SZJL201907), a project funded by the Priority Academic Program Development of Jiangsu Higher Education Institutions (PAPD), the Hui-Chun Chin and Tsung-Dao Lee Chinese Undergraduate Research Endowment (CURE).

## Author contributions

L.Z. and J.X. designed and supervised the study. J.G., Z.T., J.L., Y.C., S.L., Q.G., Y.C., and S.L. performed experiments. J.G., L.Z., and J.X. analyzed data. J.G., L.Z., and J.X. wrote the paper. All authors contributed to, read, and approved the final manuscript.

## Ethics

Animal experimentation: All of the animal studies were treated in accordance with the guidelines in the Care and Use of Animals and with the approval of the Soochow University Animal Welfare Committee (approval number: 201906A099).

